# Systematic generation and imaging of tandem repeats reveal multivalent base-pairing as a major determinant of RNA aggregation

**DOI:** 10.1101/2022.04.11.487960

**Authors:** Atagun U. Isiktas, Aziz Eshov, Suzhou Yang, Junjie U. Guo

**Affiliations:** Department of Neuroscience, Yale University School of Medicine, New Haven, CT 06520, USA; Interdepartmental Neuroscience Program, Yale University, New Haven, CT 06520, USA

**Keywords:** Short tandem repeat, repeat expansion disorder, RNA aggregation, multivalent interactions, RNA base pairing, RNA imaging

## Abstract

Expansion of short tandem repeats (STRs) in the human genome underlies over fifty genetic disorders. A common pathological feature of repeat RNAs is their propensity to aggregate in cells. While these RNA aggregates have been shown to cause toxicity by sequestering RNA-binding proteins, the molecular mechanism of repeat RNA aggregation remains unclear. Here we devised a generalizable method to efficiently generate long tandem repeat DNAs de novo and applied it to systematically determine the sequence features underlying RNA aggregation. Live-cell imaging of repeat RNAs indicated that aggregation was mainly driven by multivalent RNA-RNA interactions via either canonical or noncanonical base pairs. While multiple short runs of two consecutive base pairs were sufficient, longer runs of consecutive base pairs such as those formed by a neurodegeneration-associated hexanucleotide repeat further enhanced aggregation. In summary, our study provides a unifying model for the molecular basis of repeat RNA aggregation and a generalizable approach for identifying the sequence and structural determinants of repeat RNA properties.

## INTRODUCTION

An increasing number of human diseases, especially neurological disorders, have been shown to arise from STR expansion mutations. While some of them are recessive mutations (e.g., fragile X syndrome), many repeat expansion mutations are inherited in a dominant manner, suggesting possible involvement of gain-of-function mechanisms (Rodriguez and Todd, 2019; Schwartz et al., 2021). One such mechanism that has been well established in myotonic dystrophy type 1 and 2 (DM1/2) is mediated by RNA transcripts containing the expanded repeat sequence, which form aggregates also known as repeat RNA foci (Ranum and Cooper, 2006). The aggregated CUG and CCUG repeat RNAs in DM1 and DM2, respectively, sequester several RNA-binding proteins (RBPs) including the muscleblind-like (MBNL) family of splicing regulators, thereby causing widespread dysregulation in pre-mRNA splicing (Scotti and Swanson, 2016; Wang et al., 2012). In addition to DM1/2, repeat RNA foci have been widely observed in other repeat expansion disorders such as Huntington’s disease (CAG repeats) (Didiot et al., 2018) and *C9orf72*-associated amyotrophic lateral sclerosis (ALS) (DeJesus-Hernandez et al., 2011; Zu et al., 2013) and frontotemporal dementia (FTD) (GGGGCC repeats). However, the pathophysiological significance of repeat RNA foci in these diseases is less well understood.

By imaging CAG/CUG and GGGGCC repeat RNA foci assembly and disassembly in cells, a previous study has shown that these RNA foci are liquid-like, dynamic structures that share similarities with phase-separated protein condensates formed by multivalent interactions (Jain and Vale, 2017). The same study has further proposed that repeat RNAs may assemble by directly base-pairing with one another, a model reminiscent of cytoplasmic stress granule assembly (Van Treeck et al., 2018). While this model is plausible, direct evidence of a critical role of RNA-RNA base-pairing is currently lacking. Non-mutually exclusive, alternative mechanisms of repeat RNA aggregation may involve, for example, RBPs that contain low-complexity domains that are prone to multimerize via either homotypic or heterotypic interactions (Kato et al., 2012). Once recruited by repeat RNAs, such RBPs may facilitate condensate formation, which can in turn recruit additional repeat RNAs in an indirect manner.

For understanding the mechanism of repeat RNA aggregation, it is important to identify the sequence features that promote this process. However, previous studies of repeat RNA foci have been largely restricted to disease-associated expanded repeats, which represent only a small fraction of all possible repeat sequences. In addition, most dominantly pathogenic repeats have been shown to form RNA foci to some extent, with few counterexamples available. Moreover, each repeat region is endogenously expressed in distinct genomic context and cellular environment, thereby further complicating their direct comparison.

To circumvent these confounding factors, we developed a generalizable method to efficiently generate long tandem repeat DNA fragments of any desired sequence and length, which can be subsequently cloned in expression constructs for functional assays. By using this approach, we monitored the formation of RNA foci for a variety of repeats, most of which have not been previously studied. Results from this unbiased analysis uncovered the sequence determinants of repeat RNA aggregation and strongly supported RNA-RNA base pairing as a main cause of repeat RNA foci formation.

## RESULTS

### Generation of long tandem repeats by RepEx-PCR

To investigate the sequence determinants of repeat RNA aggregation, we needed an efficient and generalizable method to synthesize long tandem repeats of any desired sequence. While a previously reported method based on type IIS restriction endonucleases allows sequential elongation of any repeat (Scior et al., 2011) (hereafter referred to as the sequential method), it requires multiple rounds of cloning to create long repeats in the pathological range. We took inspiration from the seminal work by Khorana and colleagues on generating di- and trinucleotide repeat DNA polymers by using *E. coli* DNA polymerase I (Khorana et al., 1965), and devised a PCR-based method to generate long DNA repeats from short DNA oligos, which we termed repeat-extension PCR (RepEx-PCR) (**Figure 1A**). RepEx-PCR is initiated by annealing two complimentary single-stranded DNA oligos (typically 16-20 nt) containing two or more repeats of the desired sequence. While most oligos would form perfect duplexes, some would anneal in an offset manner, yielding either 5’ or 3’ overhangs. Next, *Taq* DNA polymerases fill in the 5’ overhangs with additional repeat units, before all duplexes are denatured for the next cycle of annealing and extension. In addition to filling 5’ overhangs, *Taq* polymerases with 3’-to-5’ proofreading exonuclease activity remove 3’ overhangs, thereby blunting both ends of the double-stranded repeat DNA products (**Figure 1A**).

**Figure 1.**
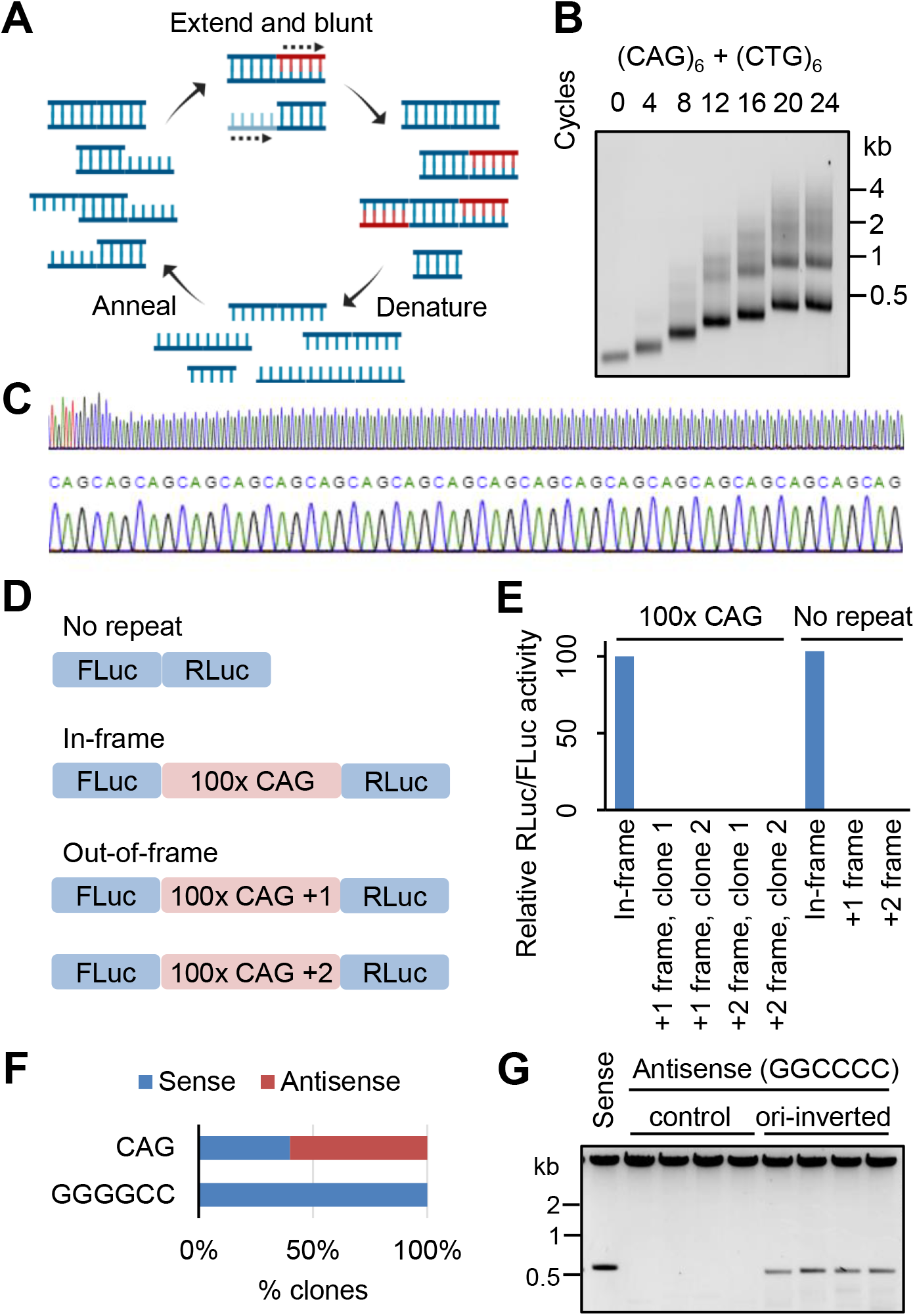
Generation of long tandem repeats by RepEx-PCR. (**A**) Schematic illustration of RepEx-PCR. (**B**) Length distribution of RepEx-PCR products. (C) Sanger sequencing results of a ~100x CAG construct, showing the junction between 5’ flanking region and repeat insert (*top*) and a magnified view of CAG repeats (*bottom*). (D) Schematic illustration of a dual-luciferase reporter for detecting frameshifted CAG repeats. (**E**) Relative RLuc/FLuc activities of out-of-frame 0x or 100x CAG constructs, normalized to the respective in-frame controls. (**F**) Percent of clones with CAG or GGGGCC repeats inserted in the sense and antisense direction. (**G**) Restriction enzyme digestion of clones containing antisense (GGCCCC) repeats inserted in the original or ori-inverted vector.

As expected, increasing numbers of RepEx-PCR cycles generated longer repeat DNAs spanning a wide range in length. For example, RepEx-PCR starting from two oligos containing six units (6x) of CAG and CTG repeats, respectively, generated CAG:CTG repeat DNA products ranging from tens to thousands of copies (**Figure 1B**), well into the pathological range associated with Huntington’s disease and myotonic dystrophy type 1. Size-selected repeat DNA products were 5’-phosphorylated, ligated into cloning vectors, amplified in *E. coli*, and verified by sequencing (**Figure 1C**). In principle, RepEx-PCR products should be consisted only of full repeats. To test whether incomplete repeats caused by small indels may arise during either RepEx-PCR or subsequent cloning, we generated a series of dual luciferase reporter constructs containing CAG repeats (**Figure 1D**). In these reporters, ~100x CAG repeats were in-frame with the upstream firefly luciferase (FLuc) coding sequencing, while a downstream Renilla luciferase (RLuc) coding sequencing was in either 0- (in-frame), +1-, or +2-frame (out-of-frame) relative to FLuc. If RepEx-PCR or subsequent cloning generated incomplete repeats, we would observe decreased RLuc activity from the in-frame reporter and increased RLuc activity from +1 or +2-frame reporters. After transfection in HEK293T cells, all four tested clones of out-of-frame reporters yielded background RLuc activity similar to the control vectors with no repeats inserted (**Figure 1E**), suggesting that most if not all RepEx-PCR products are pure tandem repeats with no incomplete repeat units.

Blunt-end ligation of RepEx-PCR products should in principle result in repeats being inserted in either the sense or antisense orientation. Indeed, we obtained similar numbers of clones containing either CAG or CTG repeats (**Figure 1E**). When we applied RepEx-PCR to generate other repeats, however, some repeats showed strong orientation bias. For example, RepEx-PCR for the hexanucleotide GGGGCC repeats only yielded clones in sense orientation but not in the antisense (GGCCCC) orientation (**Figure 1F**). Attempts to reverse the G-rich repeats by using restriction enzymes resulted in the loss of repeat inserts (**Figure 1G**), indicating that the observed orientation bias was largely due to repeat instability instead of ligation bias. Similar strand-specific repeat instability has been previously attributed to the position of repeats relative to the origin of replication (ori) (Kang et al., 1995)). Indeed, inverting the ori before ligation substantially stabilized the repeats in the antisense orientation (**Figure 1G**).

### Sequence determinants of triplet repeat RNA aggregation

An MS2 stem-loop/MS2 coat protein (MCP)-based RNA imaging system has previously been used to monitor repeat RNA aggregation in living cells (Jain and Vale, 2017) (**Figure 2A**). After transfecting 12x MS2 stem-loop-tagged repeat RNA expression constructs in U2OS cells that stably express MCP-YFP and reverse tetracycline-controlled trans-activator (rtTA), we induced repeat RNA expression by adding doxycycline and imaged the formation of RNA foci the next day. By comparing 20x and ~200x CAG repeats, we first confirmed the previous observation that repeat RNA aggregation was strongly dependent on repeat length (Jain and Vale, 2017) (**Figure 2B**). To determine the sequence features that promote CAG repeat RNA foci formation, we applied RepEx-PCR to generate variants of CAG repeats with similar length (~200x), each containing substitutions of one of the three nucleotides. Substituting C1 with G (GAG repeats) or G3 with C (CAC repeats) abolished the formation of RNA foci, whereas A2U substitution (CUG repeats) had little effect (**Figure 2C**). These results were consistent with the previously proposed model (Jain and Vale, 2017), in which CAG repeat RNAs form multivalent interactions via direct C:G base pairs. To further assess the role of base pairing, we generated the compensatory double mutant C1G/G3C (GAC repeats), in which consecutive C:G pairs were restored. Indeed, GAC repeat RNAs readily formed foci in U2OS cells (**Figure 2C**).

**Figure 2.**
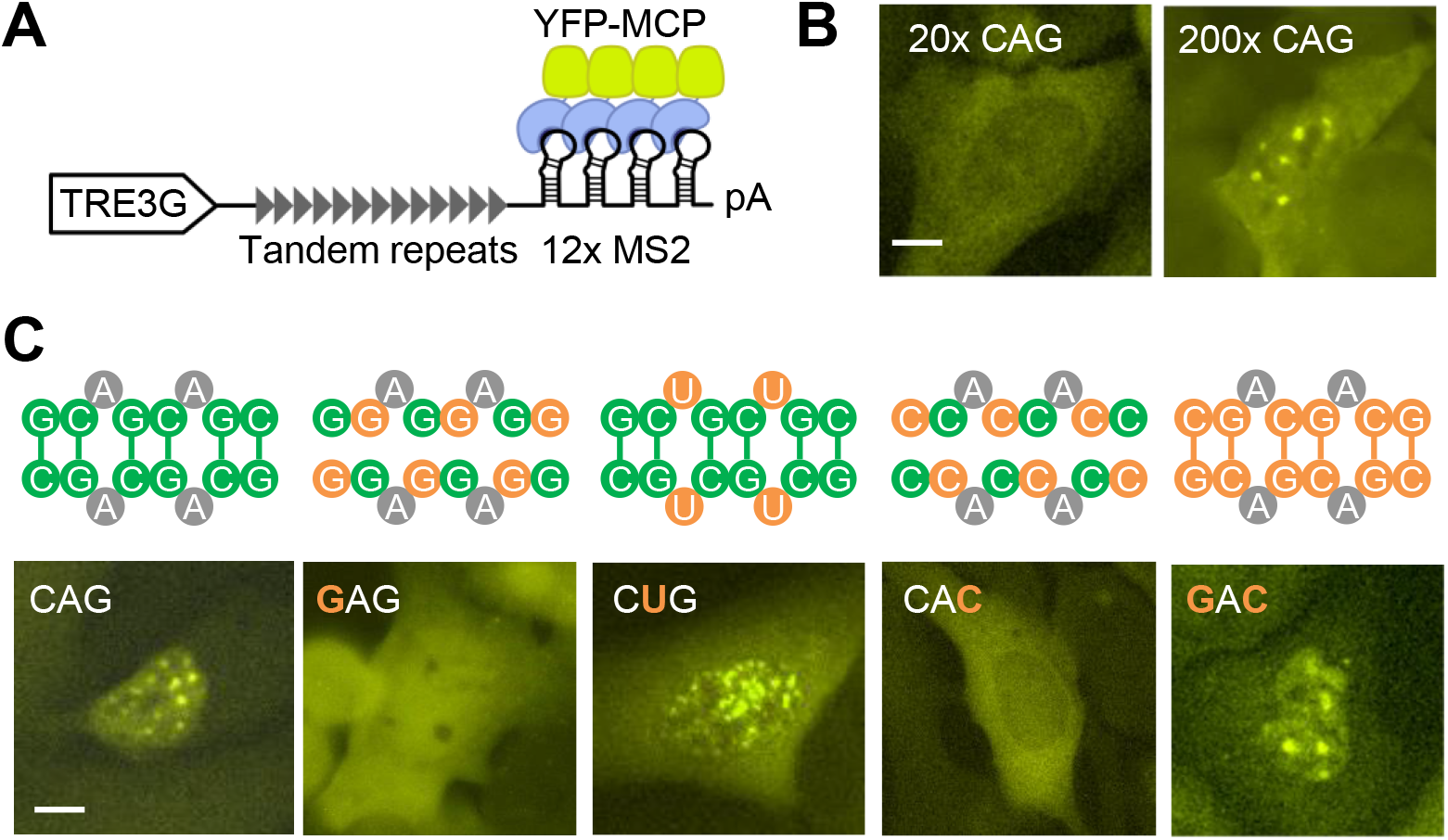
Sequence determinants of CAG repeat RNA aggregation. (**A**) Schematic illustration of the 12x MS2-tagged doxycycline-inducible repeat RNA imaging construct. (**B**) Representative images of cells expressing 20x (*left*) and ~200x (*right*) CAG repeat RNAs. Scale bar, 10 μm. (**C**) Potential base pairing patterns (*top*) and representative images (*bottom*) of CAG sequence variants. Substituted nucleotides are highlighted in orange. See Figure 3C for quantification. Scale bar, 10 μm.

To more systematically determine the sequence characteristics that promote repeat RNA aggregation, we sought to generate all possible triplet RNA repeats and quantify their aggregation in cells. Except for the four homopolymers (polyA, polyC, polyG, and polyT), we generated all of the 20 remaining non-redundant triplet repeats (**Figure 3A**). Consistent with the previous analysis on CAG variants, we restricted the length at ~600 bp or ~200x. Because the percentage of foci-positive cells was highly correlated with the average RNA foci area per cell (**Figure S1**), we used the former as our primary measurement.

**Figure 3.**
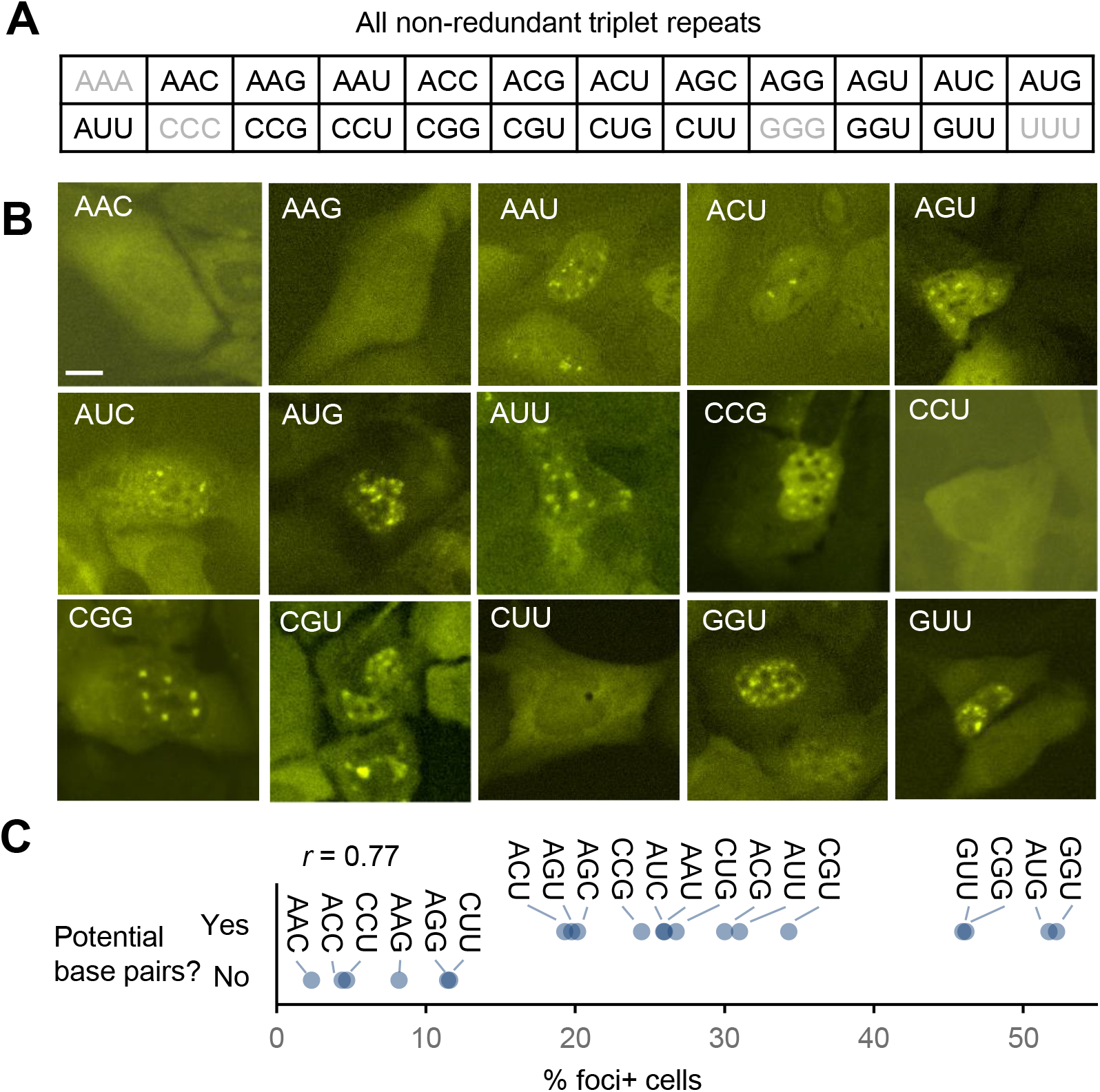
Expanded analysis of trinucleotide repeat RNA aggregation. (**A**) List of all of 20 non-redundant triplet repeats. Homopolymers are indicated in gray. (**B**) Representative images of cells expressing each of 15 triplet repeats not included in Figure 2. Scale bar, 10 μm. (**C**) Relationship between predicted base pairs and foci forming ability. Pearson’s correlation coefficient is shown.

Similar to CAG variants, the 20 tested triplet repeat RNAs exhibited a wide range of aggregation propensity (**Figure 3B**), which was not explained by differences in the expression level of each repeat sequence (**Figure S2**). To further rule out the potential effect of differential expression between repeats, we measured repeat RNA expression levels and foci formation at each of a series of doxycycline concentrations, and then interpolated the percentage of foci-positive cells at a fixed expression level (~50% of GADPH expression level measured by RT-qPCR), which yielded highly consistent results with those from experiments using a constant doxycycline concentration (**Figure S3**).

The 20 triplet repeat sequences allowed us to assess more broadly the role of base pairing in repeat RNA aggregation. Consistent with our CAG variant analysis, binary categorization of triplet repeats based on their potential to form Watson-Crick or G:U wobble base pairs was remarkably predictive of foci formation (**Figure 3C**), accounting for more than half of the variance (*r*=0.77). These results strongly supported a primary role of RNA base-pairing in mediating repeat RNA aggregation.

### Base pairing properties in repeat RNA aggregation

We next sought to further characterize the type and strength of base pairing required for repeat RNA aggregation. For triplet repeats, base pairs were invariably in runs of two (**Figure 2C**). To test whether multiple runs of two consecutive base pairs were the minimal interactions, we generated ~150x ACAG repeats by RepEx-PCR, which could potentially form non-consecutive C:G pairs (**Figure 4A**). In contrast to 200x CAG repeats, 150x ACAG repeat RNA did not form foci (**Figure 4B, C**), suggesting that runs of as few as two consecutive base pairs were the minimal requirement for RNA aggregation.

**Figure 4.**
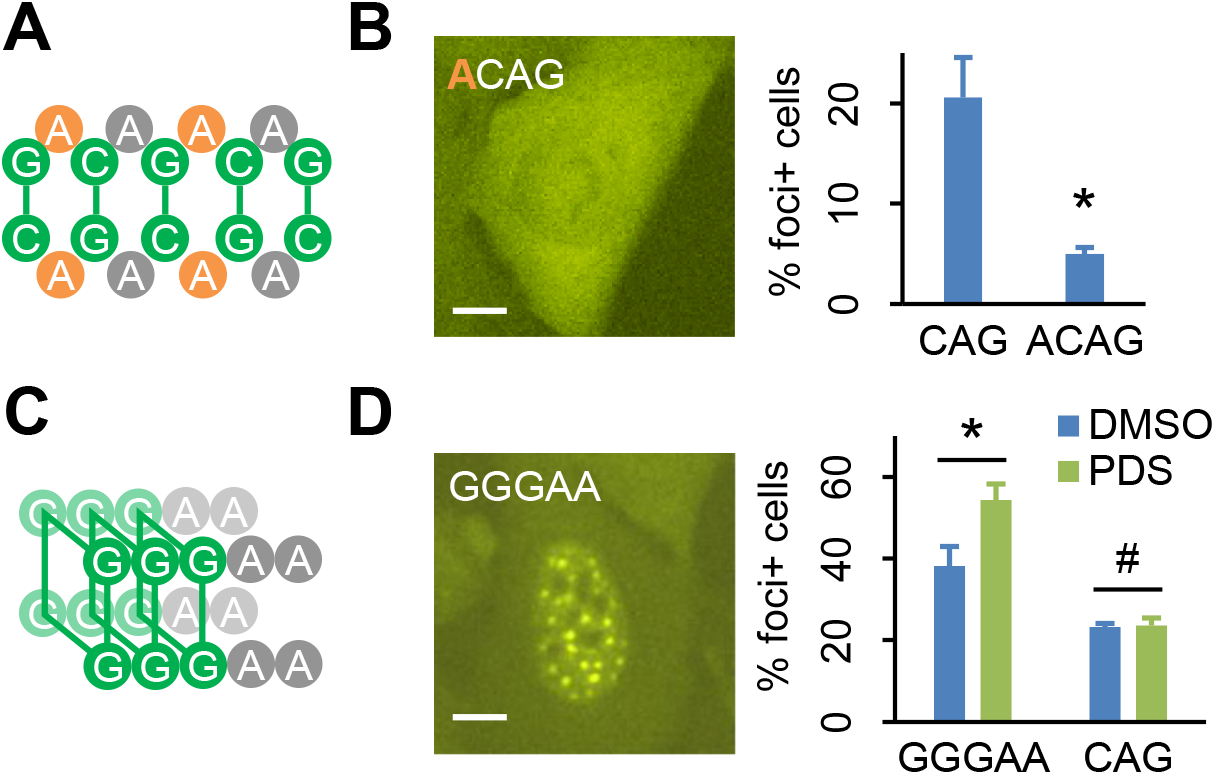
Base pairing properties in repeat RNA aggregation. (**A**) Potential base-pairing pattern of ACAG repeats. The added A nucleotides are highlighted in orange. (**B**) Representative image (left) and quantification (*right*) of cells expressing 150x ACAG repeat RNA. Scale bar, 10 μm. *, p<0.05, two-tailed Student’s t test. (**C**) Predicted G-quadruplex structure of GGGAA repeats. (**D**) Representative image (left) and quantification (*right*) of cells expressing 120x GGGAA or 200x CAG repeat RNAs, treated with DMSO or PDS. Scale bar, 10 μm. #, p>0.1; *, p<0.05, two-tailed Student’s t test.

Previous studies have proposed that repeat RNAs containing multiple consecutive G nucleotides may aggregate via the formation of G-quadruplex (Fay et al., 2017; Jain and Vale, 2017), a four-stranded RNA structure mediated by noncanonical G:G base pairs. Of the three tested triplet repeats containing GG dinucleotides, we observed RNA foci formation for CGG and UGG, but not AGG (**Figure 2C**). However, CGG and UGG foci could be due to C:G and G:U wobble base pairs, respectively, whereas the scarcity of AGG foci might be due to RBPs and helicases that normally prevented RNA G-quadruplex formation in cells (Guo and Bartel, 2016). To test whether repeat RNA aggregation could be mediated by stronger G-quadruplex interactions, we generated 120x repeats of AAGGG, a well-established G-quadruplex motif (Guo and Bartel, 2016) (**Figure 3C**). Not only 120x AAGGG repeat RNAs strongly aggregated (**Figure 3D**), treating cells with pyridostatin (PDS), a G-quadruplex stabilizer (Biffi et al., 2014), further increased AAGGG but not CAG foci (**Figure 3D**). These results suggested that aside from runs of two (or more) consecutive canonical base pairs, multiple runs of three (or more) consecutive G nucleotides could also mediate repeat RNA aggregation through noncanonical base pairs and G-quadruplex assembly.

### Aggregation of ALS/FTD-associated GGGGCC repeat RNAs

Expansion of a hexanucleotide GGGGCC repeats within the first intron of *C9orf72* gene is the most common genetic cause of amyotrophic lateral sclerosis (ALS) and frontotemporal dementia (FTD) (DeJesus-Hernandez *et al*., 2011; Renton et al., 2011). Similar to other pathogenic repeats, GGGGCC repeat as well as its antisense GGCCCC repeat RNAs are both known to form foci in patient tissues (DeJesus-Hernandez *et al*., 2011; McEachin et al., 2020; Zu *et al*., 2013). In keeping with the notion that runs of consecutive base pairs mediates RNA aggregation, GGGGCC repeats could form runs of four consecutive base pairs in multiple configurations (**Figure 5A**), two of which involving canonical C:G pairs between C5/C6 and either G3/G4 (*a*) or G1/G2 (*b*). GGGGCC repeats could also form four-layer G-quadruplexes (*c*), as have been previously shown in vitro (Fay *et al*., 2017; Fratta et al., 2012; Jain and Vale, 2017; Reddy et al., 2013), as well as a variety of configurations involving shorter runs of base pairs. Consistent with previous studies, we observed strong aggregation propensity of GGGGCC repeat RNAs in a repeat length-dependent manner (**Figure 5B**), with 60-80% transfected cells showing clear RNA foci upon induction. To test whether the strong base-pairing potential of GGGGCC repeat RNA may promote foci formation, we generated three mutant sequences, each containing substitutions that disrupted two of the three possible “run-of-four” configurations (**Figure 5C**). Specifically, AGGGCC (G1A) repeats could still form configuration *a*, as well as a three-layer G-quadruplex (not shown), whereas GGAGCC (G3A) and GGGGAC (C5A) repeats could form configurations *b* and *c*, respectively. Indeed, with the remaining ability to form runs of four consecutive base pairs, each of the three mutants still strongly aggregated (**Figure 5C, D**). In contrast, foci formation of a G1A/G3A double mutant, which could only form runs of two consecutive C:G pairs (**Figure 5C**), was significantly decreased to a level comparable to other GC-rich triplet repeats (**Figure 5C, D**). These results suggested that longer runs of consecutive base pairs enhanced GGGGCC repeat RNA aggregation, further supporting the role of base-pairing in promoting repeat RNA foci formation.

**Figure 5.**
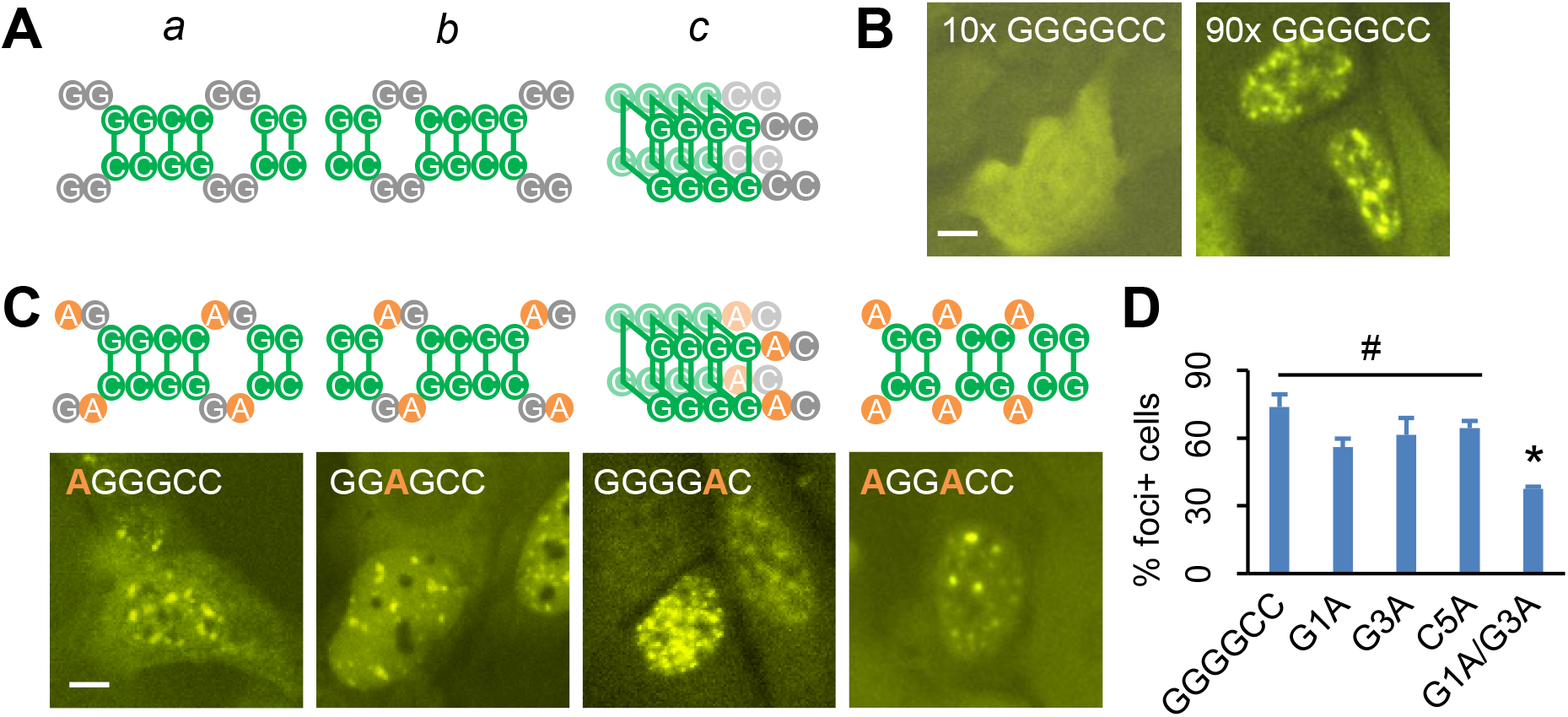
Aggregation of ALS/FTD-associated GGGGCC repeat RNAs. (**A**) Potential base-pairing patterns of GGGGCC repeats. (**B**) Representative images of cells expressing 10x (*left*) or 90x (*right*) GGGGCC repeat RNAs. Scale bar, 10 μm. (**C**) Potential base-pairing patterns (*top*) and representative images (*bottom*) of cells expressing each GGGGCC variant. Substituted nucleotides are highlighted in orange. Scale bar, 10 μm. (**D**) Quantification of foci+ cells expressing each GGGGCC variant. #, p>0.1; *, p<0.05, two-tailed Student’s t test

## DISCUSSION

Our PCR-based repeat extension provides an efficient and generalizable approach to generate tandem repeats of any desired sequence spanning a wide range of lengths. Compared to a previous type IIS restriction endonuclease-based method (the sequential method) (Scior *et al*., 2011), RepEx-PCR has two clear advantages: First, the sequential method relies on a pair of type IIS restriction sites flanking the repeats, which may interfere with certain applications. In contrast, blunt-end ligation of RepEx-PCR products do not require any specific flanking sequence. Second, the sequential method starts with a relatively short synthetic repeat and doubles the repeat length after each round of cloning. Therefore, multiple rounds of cloning are needed to obtain more than a few dozen repeats. On the contrary, RepEx-PCR can generate long repeats in a single round of cloning, with the minor trade-off being that multiple bacterial colonies need to be screened in order to identify those with the desired length and orientation. Nonetheless, the sequential method, due to its PCR-free nature, provides a complementary strategy that may be more suited for assembling multiple repeat sequences in the same construct.

RepEx-PCR enabled us to generate a large variety of repeat RNAs and deduce the sequence features that promote RNA foci formation. First, mutation analysis of CAG repeats showed disrupting the predicted C:G pairs formed between C1 and G3 abolished RNA aggregation (**Figure 2**), whereas the compensatory double mutations that restored base pairing also restored RNA aggregation. These observations were corroborated by our expanded analysis that surveyed all 20 non-redundant triplet repeats (**Figure 3**), in which binary categorization of base-pairing potential could effectively predict foci formation, accounting for more than half of the variance. Finally, to test whether runs of consecutive base pairs were required for RNA aggregation, we interrupted the GC and GGCC/CCGG motifs in CAG (**Figure 4**) and GGGGCC repeats (**Figure 5**), respectively, both causing significant reductions in RNA foci. Collectively, these results supported and expanded the previously proposed model in which RNA-RNA base pairs serve as the primary interactions mediating repeat RNA aggregation (Jain and Vale, 2017).

Our analysis also revealed important contributions of noncanonical RNA-RNA interactions in repeat RNA aggregation. One of the most stable noncanonical RNA-RNA interactions is the G-quadruplex structure, in which adjacent runs of consecutive G nucleotides pair to two other G-runs with both Watson-Crick and Hoogsteen faces. While many endogenous RNAs contains regions that can form G-quadruplexes in vitro, these regions are overwhelmingly unfolded in eukaryotic cells by RNA helicases and RBPs (Guo and Bartel, 2016), presumably to prevent their potential negative impact on translation and degradation. In contrast to physiological mRNAs, some repeat RNAs such as those in *C9orf72*-associated ALS/FTD and SCA36 (UGGGCC repeats) contain hundreds of adjacent G-runs and therefore have extraordinarily high propensity to form intra- or intermolecular G-quadruplexes (Fratta *et al*., 2012; Reddy *et al*., 2013). Indeed, our analyses of GGGAA and GGGGCC repeats provided evidence that expanded repeat RNAs with runs of three or more G nucleotides could readily aggregate via G-quadruplexes, suggesting that unlike transiently formed G-quadruplexes in physiological RNAs, G-quadruplexes within repeat RNA aggregates might become less accessible to the cellular remodeling machinery (Conlon et al., 2016). These results also raised the intriguing possibility that G-rich repeat RNA foci might sequester G-quadruplex-unfolding factors and thereby stabilize G-quadruplexes within physiological RNAs in trans.

On one hand, our finding that multiple runs of two or more consecutive base pairs were sufficient to promote RNA aggregation explains why RNAs containing expanded repeats, with their high density of base-pairing motifs, were particularly prone to aggregation. On the other hand, this model also suggests that RNA aggregates may not be consisted purely of RNAs from the expanded repeat locus but also of other endogenous RNAs with the potential to base-pair extensively to repeat RNAs, such as pre-mRNAs with similar albeit non-expanded STR sequences, potentially causing the misprocessing or retention of these secondarily recruited RNAs. Considering that previous studies have mostly focused on the RBPs sequestered within repeat RNA foci, we anticipate that investigating the secondary RNA components may shed new light on the pathophysiological significance of repeat RNA aggregates.

While the base pairing potential explained more than half of the variance in RNA foci formation between repeat sequences, some sequence features associated with foci formation were not readily explained by base-pairing. For example, while G:U wobble pairs are in general weaker than C:G pairs, GGU and GUU repeats were more prone to aggregation than GGC and GCC repeats, respectively (**Figure 3C**), suggesting that G/U nucleotides may promote RNA aggregation in ways additional to base pairing, possibly due to their interactions with certain RBPs. Our method and results should enable future studies in identifying RBPs and RNA helicases that influence repeat RNA aggregation. Finally, our systematic approach of identifying sequence and structural features in repeat RNAs is not limited to the studies of RNA foci, but generalizable for studying other properties of repeat sequences, such as their impact on pre-mRNA processing (Sznajder et al., 2018), non-AUG translation (Guo et al., 2022), antisense RNA expression (Zu *et al*., 2013), cytoplasmic stress granules (Fay *et al*., 2017), and cell viability (Sun et al., 2020). Therefore, RepEx-PCR represents a valuable addition to the repertoire of methods enabling deeper investigations into the physiology and pathophysiology of tandem repeats and repeat expansion disorders.

## METHODS

### Generation of tandem repeats by RepEx-PCR

RepEx-PCR was performed by using 0.5 μM each of the two single-stranded DNA oligos (Integrated DNA Technologies) containing 5-7x sense and antisense repeats, respectively. For most triplet repeats, Q5 hot-start high-fidelity DNA polymerase (New England Biolabs) was used. For GC-rich repeats such as GGGGCC, KAPA HiFi DNA polymerase (Roche) was used with GC buffer and/or GC enhancer. RepEx-PCR products were size-selected and purified from a 0.8% agarose gel, 5’ phosphorylated by T4 polynucleotide kinase (New England Biolabs), ligated to a blunt-end linear vector (typically also PCR products), and transformed into NEB Stable competent cells (New England Biolabs). After overnight incubation, individual bacterial colonies were amplified in liquid cultures before plasmid DNA was extracted. Repeat insert size and orientation were determined by restriction enzyme digestion and DNA sequencing, respectively. After sequencing validation, the repeat insert was further subcloned to a doxycycline-inducible (Tet-On, Clontech) expression vector containing 12x MS2 hairpins.

### Luciferase assays

RepEx-PCR-generated ~100x CAG repeats were inserted into a dual-luciferase reporter plasmid modified from pmirGLO (Promega) between firefly luciferase (FLuc) coding sequence in the 0 frame and Renilla luciferase (RLuc) coding sequence in the 0, +1, or +2 frame. Reporter plasmid DNAs as well as control reporter plasmid DNAs with no repeats inserted were transfected in HEK293T cells in 24-well plates using Lipofectamine 2000 (ThermoFisher). 16-24 hours after transfection, cells were lysed in Glo Lysis Buffer (Promega) at room temperature for 5 min. FLuc and RLuc activities in lysates were sequentially measured by using Dual-Glo luciferase assay reagents and a GloMax 20/20 luminometer (Promega) according to the manufacturer’s instructions.

### Live-cell imaging of repeat RNA foci

The U2OS cell line stably expressing rtTA and MCP-YFP was a gift from Jain and Vale (Jain and Vale, 2017). Cells were cultured at 37°C and 5% CO_2_ in Dulbecco’s Modified Eagle Medium (DMEM) (Thermo Fisher) supplemented with 10% fetal bovine serum (FBS) (Thermo Fisher). U2OS cells plated in glass-bottom 24-well plates (Cellvis) were transfected with 320 ng 12x MS2-tagged repeat expression construct and 80 ng CMV-BFP plasmid DNA (for labeling transfected cells) mixed with 1.6 μl FuGENE HD (Promega) according to the manufacturer’s instructions. 4-6 hours after transfection, the medium was replaced with fresh DMEM supplemented with 10% FBS and 1 μg/ml doxycycline (Sigma). 16-24 hours after induction, cells were imaged at 37 °C by using a Lionheart FX automated microscope (Agilent) with 20x and 60x objectives. Typically, 50-100 BFP-positive cells were imaged in each well for RNA foci quantification.

### RT-qPCR

After live-cell imaging, total RNA from U2OS cells were extracted by using Monarch Total RNA Miniprep kit (New England Biolabs) and treated with TURBO DNase (Thermo Fisher). RT-qPCR was performed by using Luna Universal One-Step RT-qPCR reagents (New England Biolabs). Primers targeting the MS2 stem-loop regions were used for quantifying repeat RNA expression levels, which were normalized by GAPDH expression levels.

## ACKNOWLEDGEMENT

We thank D. Bartel for the initial suggestion of using PCR for repeat synthesis and members of the Guo lab for discussions and comments on the manuscript. This work is supported by an NIH New Innovator Award (DP2 GM132930) and the Muscular Dystrophy Association (MDA602934). J.U.G. is a Klingenstein–Simons Fellow in Neuroscience and a New York Stem Cell Foundation–Robertson Investigator.

---

**Figure S1.**
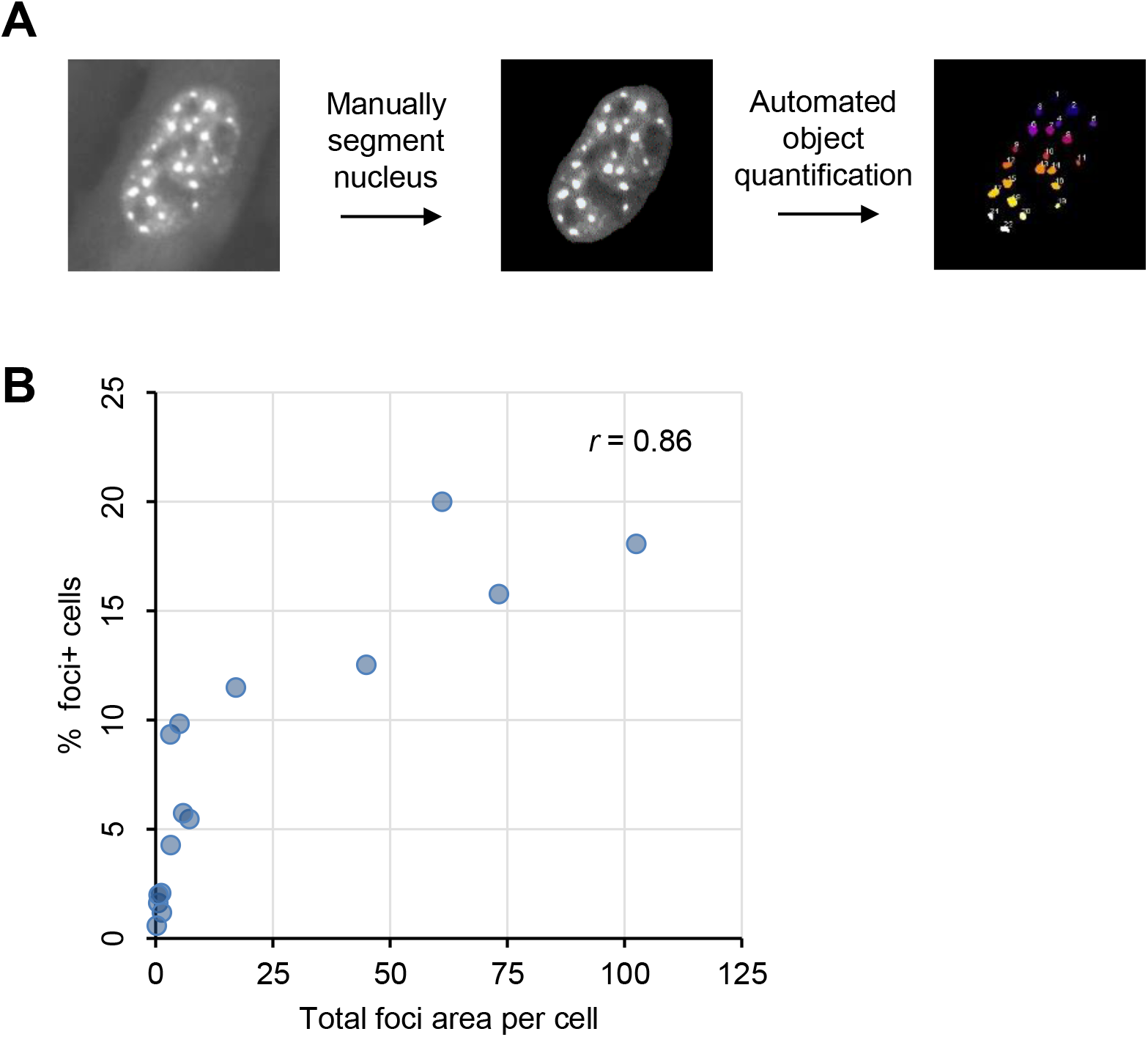
Relationship between the percentage of foci+ cells and the size of RNA foci per cell. (**A**) Procedure of RNA foci quantification. We used FIJI to analyze the acquired images, manually segmented the nucleus of each cell, and quantified the mean YFP fluorescence intensity in each nucleus. Next, we used the 3D Objects Counter plugin with an intensity threshold of 1.6x the mean YFP fluorescence intensity and a size threshold of 15 adjacent pixels. (**B**) Correlation between the percentage of foci+ cells and total foci area per cell for each triplet repeat. Pearson’s correlation coefficient is shown.

**Figure S2.**
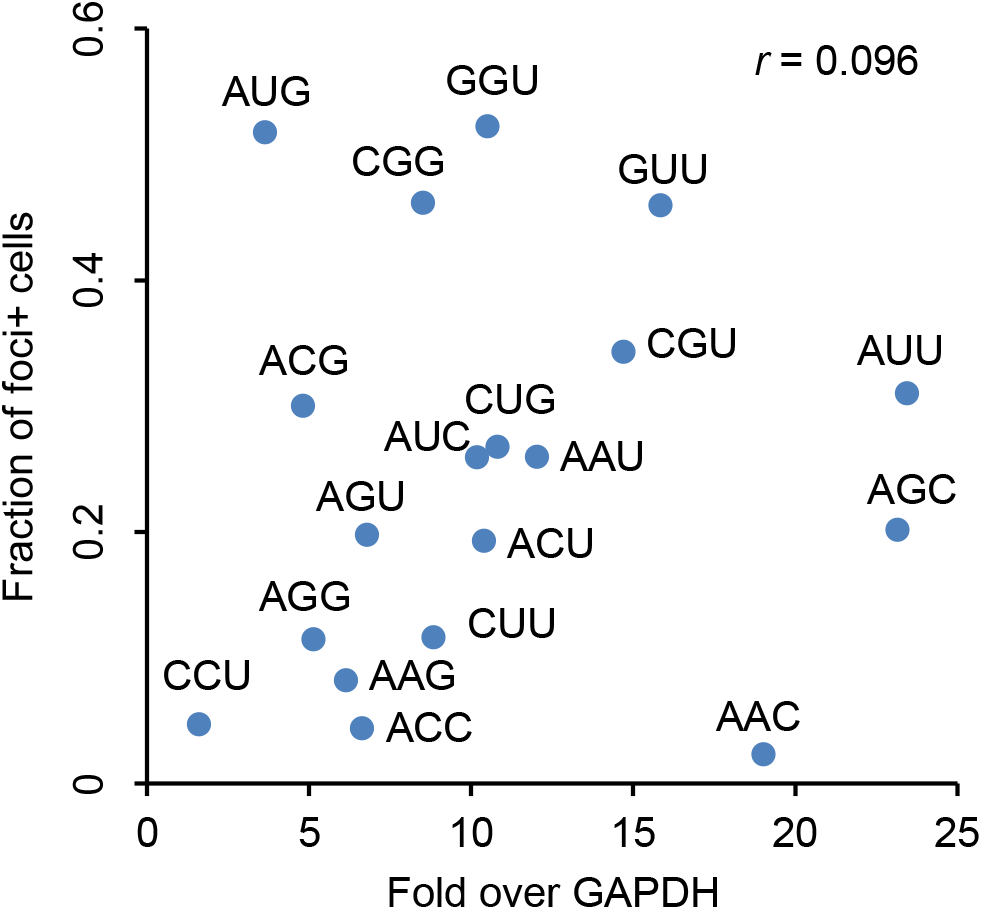
Relationship between repeat RNA abundance and foci formation. Scatter plot showing a lack of correlation between foci forming potential and expression level of each triplet repeat. Pearson’s correlation coefficient is shown.

**Figure S3.**
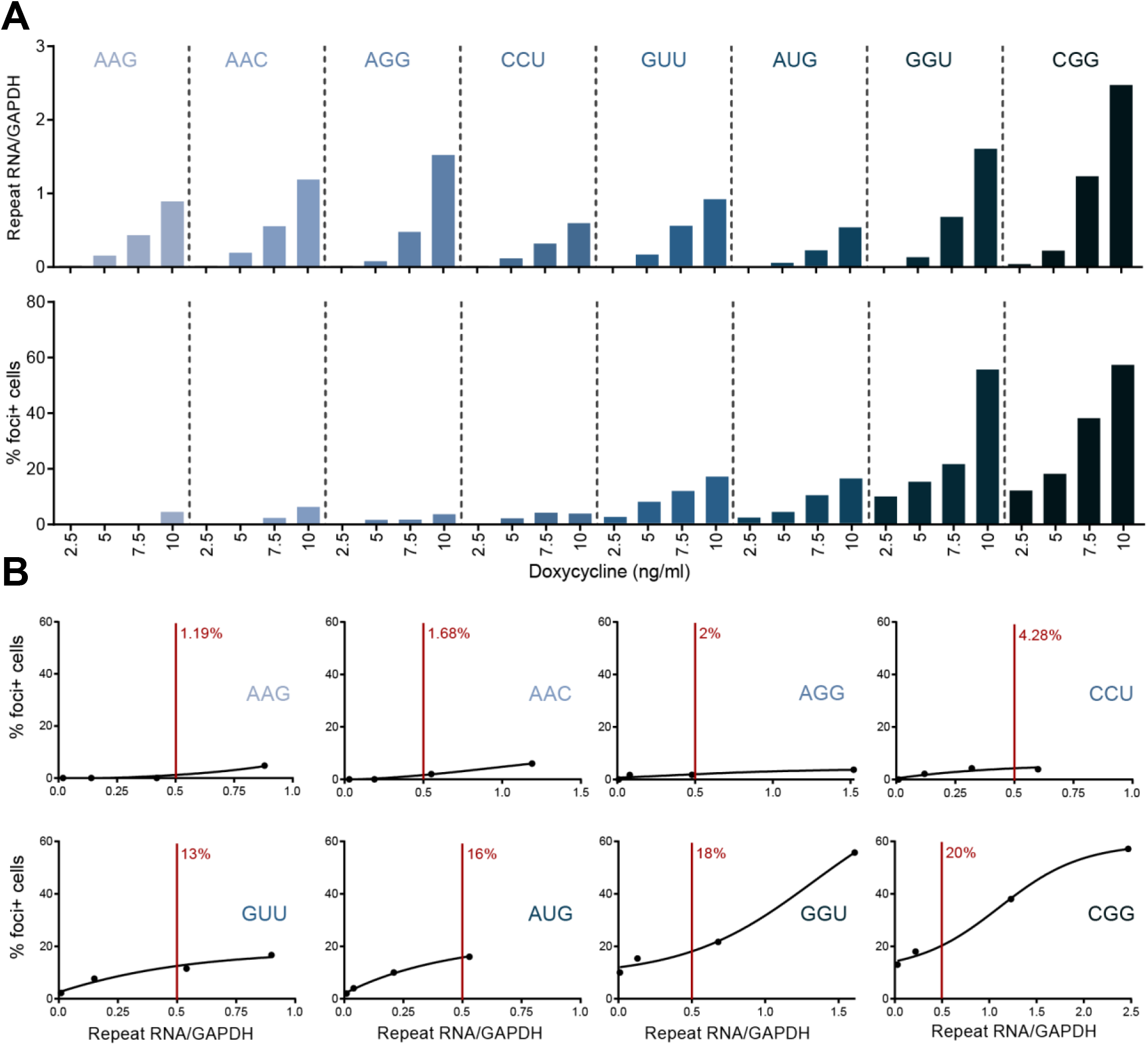
Repeat RNA aggregation at lower expression levels. (**A**) Expression levels (*top*) and percentage of foci+ cells (*bottom*) for each of eight triplet repeats at each of four doxycycline concentrations. (**B**) Dose-response relationship of RNA foci formation at increasing expression levels. Data were fit by sigmoidal functions, which were then used for interpolating the percentage of foci+ cells at a fixed expression level (0.5 fold over GAPDH).

